# Novel QTLs for cucumber resistance to two *Colletotrichum orbiculare* strains of different pathogenic races

**DOI:** 10.1101/2022.02.28.482428

**Authors:** Hiroki Matsuo, Sachiko Isobe, Kenta Shirasawa, Yosuke Yoshioka

## Abstract

*Colletotrichum orbiculare* is a hemibiotrophic fungal pathogen that causes anthracnose disease in cucumber (*Cucumis sativus* L.) and other cucurbit crops. The cucumber accession Ban Kyuri (G100) has a high level of resistance to anthracnose and is considered to be promising breeding material. Here, we report genetic mapping of loci from this cultivar that confer resistance to 104-T and CcM-1 of *C. orbiculare* that belong to pathogenic races 0 and 1, respectively. Quantitative trait locus (QTL) analysis based on phenotypic data from 196 F_2:3_ families detected one major QTL, *An5*, and one minor QTL, *An6*.*2*, for resistance to race 0, and one major QTL, *An2*, and three minor QTLs (*An1*.*1, An1*.*2* and *An6*.*1*) for resistance to race 1. We identified *lysM domain receptor-like kinase 3* (*CsaV3_5G036150*) and *wall-associated receptor kinase-like* (*CsaV3_6G048820*) as candidate genes for *An5* and *An6*.*2*, respectively. Multiple genes encoding pattern recognition receptors were located in the regions of the QTLs conferring resistance to race 1. Thus, we identified potential sources of genetic resistance to different pathogenic races of *C. orbiculare* in the Ban Kyuri cultivar of cucumber.

**Key message:** Quantitative trait locus analysis identified independent novel loci in cucumber responsible for resistance to races 0 and 1 of the anthracnose fungal pathogen *Colletotrichum orbiculare*.

## Introduction

Anthracnose is one of the most economically important foliar diseases in the production of cucumber (*Cucumis sativus* L.) and other cucurbit crops in warm and humid regions. This disease is caused by the fungal pathogen *Colletotrichum orbiculare* (Berk. et Mont.) Arx [syn. *C. lagenarium* (Pass.) Ellis et Halst.], which infects all aboveground organs such as leaves, stems, and fruits. The fungus can survive on plant residues in the field, becoming a source of infection in the next planting. The pathogen can also be transmitted through seeds. The spores and mycelial fragments of *C. orbiculare* depend upon water droplets such as rain and irrigation spray to spread. Thus, growing in greenhouses or under rain shelter is the best way to prevent anthracnose infection of cucumber crops. Open-field cultivation is relatively cost-effective, but plants are more prone to the disease than those in greenhouses. Therefore, open-field cultivation requires stricter cultivation management, such as removing plant residue from the field, improving airflow to reduce humidity, and spraying with fungicides, to suppress the spread of this disease. However, these disease control methods incur costs in time, labor, and materials, and require training and experience regarding the optimal frequency and timing of the operations. Moreover, it is difficult to control this disease completely in the rainy season, which provides the most desirable environment for anthracnose. For these reasons, the most effective way to control anthracnose is to introduce resistant cultivars.

Understanding the pathogenicity of pathogens is essential for identifying components of host resistance and breeding resistant cultivars. In the United States, seven races (race 1, 2, 3, 4, 5, 6 and 7) of *C. orbiculare* have been reported, of which races 1 and 2 are the most common (Goode 1956; Jenkins et al. 1964; Sitterly 1973; Wasilwa et al. 1993). The accession PI 197087 and its derivatives have long been known to be resistant to anthracnose (Barnes and Epps 1952; Sitterly 1973; Wyszogrodzka et al. 1987; Linde et al. 1990), and their resistance has been used widely in breeding for nearly 60 years in the United States. Recently, Pan et al. (2018) reported that *STAYGREEN* (*CsSGR*) is a candidate gene for the race 1 anthracnose resistance locus *cla* in the inbred cucumber line Gy14 derived from PI 197087. This recessive gene (hereafter referred to as *Cssgr*) carries a loss-of-susceptibility mutation and represses chlorosis under *C. orbiculare* infection. *Cssgr* also confers durable broad-spectrum resistance to downy mildew (DM, *Pseudoperonospora cubensis*) and angular leaf spot (*Pseudomonas syringae* pv. *lachrymans*) (Wang et al. 2019). However, some cultivars that were homozygous for *Cssgr* were severely damaged by highly virulent strains of *C. orbiculare* distributed in Japan (Matsuo et al. 2022); therefore, *Cssgr* does not confer resistance to these strains. Our previous study identified a novel resistant accession (Ban Kyuri) that is highly resistant to strain 104-T and moderately resistant to strain CcM-1 of *C. orbiculare* (Matsuo et al. 2022). This accession is homozygous for the wild-type allele of *CsSGR*, and its resistance levels are higher than those of accessions harboring *Cssgr*. Therefore, we hypothesized that this accession harbors novel resistance genes or a novel allele of *CsSGR* that confers anthracnose resistance.

Resistance based on adaptive and innate immunity in plants is mainly associated with pattern-triggered immunity (PTI) and effector-triggered immunity (ETI). Pattern recognition receptors (PRRs) recognize pathogen-/microbial-associated molecular patterns (PAMPs/MAMPs) and effectors and trigger PTI and ETI, respectively. PRRs are leucine-rich repeat (LRR) domain-containing proteins and include receptor-like proteins/kinases (RLPs/RLKs) and nucleotide-binding leucine-rich-repeat receptors (NLRs). Due to the repetitive motifs and rich polymorphism of LRR domain, LRRs can readily evolve new variants that bind diverse extracellular molecules (Kobe and Kajava 2001). By recognizing patterns such as PAMPs/MAMPs and effectors on pathogens, LRR domain-containing proteins can rapidly activate immune responses, such as hypersensitive response (HR) (Dangl and Jones 2001; Jones and Dangl 2006; Jones et al. 2016). RLPs/RLKs generally control PTI, which is crucial for basal resistance, and NLRs control ETI conferring cultivar-specific resistance, RLPs/RLKs and NLRs are often encoded by resistance (*R*) genes (Boutrot and Zipfel 2017; Kourelis and van Der Hoorn 2018). In addition, *R* genes are usually dominant (but sometimes recessive) genes that showed full or partial resistance to one or more pathogens (Kourelis and Van Der Hoorn 2018).

Plant genomes encode many RLPs/RLKs and NLRs that confer resistance to a wide variety of plant pathogens (Kourelis and van der Hoorn 2018). Although most plant genomes encode hundreds of predicted NLR genes, the cucumber genome was found to encode only 67–70; a similar tendency was also found in the genomes of other cucurbits (Wan et al. 2013; Yang et al. 2013; Morata and Puigdomènech 2017). Some NLRs that confer cultivar-specific resistance have been identified in cucumber: e.g., the gene for resistance to target leaf spot (TLS, *Corynespora cassiicola*), *cca-3* (Wen et al. 2015; Wang et al. 2020); the gene for resistance to scab (*Cladosporium cucumerinum*), *ccu* (Kang et al. 2011); and the gene for resistance to fusarium wilt (FOC, *Fusarium oxysporum* f. sp. *cucumerinum*), *Foc* (Zhang et al. 2014). However, cucumber seems to rely more heavily on other types of cultivar-specific resistance than other plant species. Interestingly, the cucumber genome encodes 178–192 RLKs and 42–56 RLPs; some of these genes are co-localized in the QTL region for DM resistance (Wang et al. 2014), suggesting that the genes for DM resistance may encode RLPs/ RLKs, which confer cultivar-specific resistance, similar to NLRs. Furthermore, an RLK (*CsLRK10L2*) was identified as a candidate gene for *dm4*.*1*.*2*, a sub-QTL for resistance to DM in PI 197088 (Berg et al. 2020), and a putative LRR receptor-like serine/threonine-protein kinase (*RPK2, CsGy5G015660*) was identified as a causal gene for *pm5*.*2*, a QTL for resistance to powdery mildew (PM, *Podosphaera xanthii*) (Zhang et al. 2021).

*Colletotrichum* spp. are excellent models for studying the differentiation of infection structures and the molecular basis of fungal–plant interactions (Perfect et al. 1999; O’Connell and Panstruga 2006). The *C. orbiculare* strain 104-T from Japan has been used in comprehensive studies combining genome-/molecular-based analyses to provide insights into pathogenicity, basal resistance, and non-host resistance (Kubo and Takano 2013; Kubo et al. 2016; Kubo 2018). Although the molecular mechanisms underlying its pathogenicity have already been studied, the races of *C. orbiculare* distributed in Japan had not been reported until recently. Our previous work suggested that races 1 and 2 are present in Japan. In addition, our previous work revealed that strains assigned to race 0, which infect cucumber but not watermelon, are also distributed in Japan. Ban Kyuri is highly resistant to races 0 and 2 and moderately resistant to race 1. Therefore, this accession is a promising new source of resistance to diverse anthracnose strains of different races. Here, we report that analysis of quantitative trait loci (QTLs) associated with anthracnose resistance in Ban Kyuri revealed novel QTLs that confer resistance to 104-T (race 0) and CcM-1 (race 1) independently. We discuss the location of these QTLs and those detected in previous studies as well as the prospects for using Ban Kyuri to breed new cucumber cultivars with anthracnose resistance.

## Materials and Methods

### Plant materials and mapping population

The resistant accession Ban Kyuri (G100, JP 82352) and the susceptible accession B1, a Beit Alpha-type fixed line preserved at the University of Tsukuba, were used as parental lines. We generated F_1_ progenies by reciprocal crossing between G100 and B1 [F_1_A: B1 (♀) × G100 (♂), F_1_B: G100 (♀) × B1 (♂)]. F_2_ and F_2:3_ populations were obtained by selfing one F_1_A and 196 F_2_ individuals, respectively.

### Fungal materials and preparation of inoculum

Strains 104-T (MAFF 240422; race 0) and CcM-1 (MAFF 306737; race 1) were cultured on potato dextrose agar (PDA; Nissui Pharmaceutical Co., Ltd., Tokyo, Japan) at 25 °C under light. Mycelial plugs with conidia were soaked overnight in 10% glycerol (Fujifilm Wako Pure Chemical Corporation, Osaka, Japan) and then stored at −80 °C. For inoculum preparation, a mycelial plug was transferred onto PDA plates and incubated at 25 °C under light for 10 days. The mycelium and spores were then suspended in deionized water and spread on other PDA plates. These plates were incubated at 25 °C under light for 5– 7 days. Spores formed on the surface were suspended in deionized water and filtered through a Kimwipe tissue (Nippon Paper Crecia Co., Ltd., Tokyo, Japan) to remove mycelia. Spores in suspension were counted with a hemocytometer. The concentration was adjusted to 6.0 × 10^6^ spores/mL (104-T), and 1.5 × 10^6^ spores/mL (CcM-1). Tween-20 (0.5 μL/mL; Kanto Chemical Co., Inc., Tokyo, Japan) was added to the inoculum.

### Artificial inoculation assay to evaluate the segregating population

Host response of F_2:3_ families to 104-T and CcM-1 was evaluated by cotyledon and seedling assay. A cotyledon assay was conducted to evaluate the anthracnose resistance of the 196 F_2:3_ families. G100, B1, F_1_A and F_1_B were also evaluated. Eighteen individuals per parent (B1 and G100), F_1_A, F_1_B, and F_2:3_ line were tested in two experimental replications. For germination, cucumber seeds were placed in Petri dishes with wet filter paper (Toyo Roshi Kaisha, Ltd., Tokyo, Japan) and incubated at 28 °C in the dark. Germinated seeds were transplanted into a 38-mm-diameter plastic cell tray filled with wet culture soil (Nihon Hiryo Co., Ltd., Fujioka, Japan) and incubated at 28 °C with a 12-h photoperiod in a climate chamber (Nippon Medical and Chemical Instruments Co., Ltd., Osaka, Japan). After cotyledon unfolding, the temperature in the chamber was set to 25 °C for eight days. Plants in plastic cell trays were put in plastic boxes (60 plants per box) (Akasaka Co., Ltd., Atsugi, Japan) and the inoculum (104-T or CcM-1) was sprayed on the surface of each cotyledon (ca. 0.2 mL) with a spray bottle. The boxes were then closed to maintain 100% relative humidity and kept in the dark for 24 h. The lids were opened and the plants were grown at 25 °C with a 12-h photoperiod in the climate chamber. The resistance of each inoculated plant was scored one week after inoculation. Infection severity (IS) levels were scored as follows: 0, no lesions; 1, a few (cotyledon: <5; true leaf: <25) small lesions (<3 mm in diameter); 2, several (cotyledon: >6; true leaf: >26) small lesions or a few medium lesions (3–5 mm); 3, many (cotyledon: >10; true leaf: >50) small lesions and a few to several medium lesions, or a few large lesions (>5 mm), development of chlorotic lesions; 4, several medium lesions and a few large lesions, developed chlorotic boundaries; 5, many large lesions, covering up to/more than half of the leaf; leaf is beginning to wither; 6, the leaf is wholly withered; 7, the seedling is dead. The seedling assay evaluated the disease symptoms on true leaves of the 16 F_2:3_ lines with the highest resistance or susceptibility to 104-T and CcM-1 in the cotyledon assay. Twenty individuals per parent (B1 and G100), F_1_A, F_1_B, and F_2:3_ line were tested in a trial.

### Genotyping by double-digest restriction-site-associated DNA sequencing (ddRAD-seq)

Genomic DNA was extracted from 406 F_2_ individuals and one individual of each parent (B1 and G100) using Plant Genome DNA Extraction Mini Kits (Favorgen Biotech Corp., Changzi, Taiwan). DNA samples were diluted to a final concentration of 25 μL. Libraries were constructed and sequenced as in Shirasawa et al. (2016) with minor modifications. To construct the ddRAD-seq libraries, DNA was digested with *Pst*I and *Msp*I (Thermo Fisher Scientific, Waltham, MA, USA). Digested DNA was ligated to adapters using T4 DNA ligase (Promega, Madison, WI, USA) and purified using Agencourt AMPure XP (Beckman Coulter, Brea, CA, USA) to eliminate short (<300 bp) DNA fragments. Purified DNA was amplified by PCR with indexed primers. Amplified DNA fragments were pooled and 300–900-bp fragments were fractionated on a BluePippin 1.5% agarose cassette (Saga Science, Beverly, MA, USA). These libraries were sequenced on a HiSeq4000 (Illumina Inc., San Diego, CA, USA) in 100-bp pair-end reads by the Macrogen sequencing service (Macrogen Inc., Seoul, South Korea).

### NGS data acquisition and SNP calling

Low-quality reads were removed on the basis of a quality check by PRINSEQ (Schmieder and Edwards 2011), and the sequencing data were trimmed and filtered in fastx_clipper (http://hannonlab.cshl.edu/fastx_toolkit). The data were mapped to the cucumber (Chinese Long) v3 reference sequence (Li et al. 2019) in the Cucurbit Genomics Database (Zheng et al. 2019, CuGenDB: http://cucurbitgenomics.org/) using Bowtie2 (Langmead and Salzberg 2012). The sequence alignment/map (SAM) format files were converted to binary sequence alignment/map (BAM) format files. SNP calling and genotyping were performed with the mpileup tool in SAMtools v. 0.1.15 (Li et al. 2009a) and BCFtools v. 1.9 software (Li 2011). SNPs with >5 reads in >50% of progenies were used for analysis.

### Linkage map construction and QTL analysis

The genetic linkage map of the 406 F_2_ progeny derived from the G100 × B1 cross was constructed in R/ASMap (https://cran.r-project.org/package=ASMap, Taylor and Butler 2017) and R/qtl (http://www.rqtl.org/, Arends et al. 2010) from the SNP markers obtained by ddRAD-seq analysis. The SNP markers were selected by removing low-quality loci (>80% progeny with missing values at the locus; mapped to Chr. 0). F_2_ individuals with >80% SNP markers were selected. Then *p*-values were calculated to test for deviations of genotype frequencies at each locus from the expected ratio of 1:2:1, and after Bonferroni correction for multiple testing, markers that showed significant distortion at the 5% level were removed. Genotype error rate and genotype error logarithm of the odds (LOD) scores were calculated and SNP genotypes with high genotype-error LOD score (>0) were treated as missing data. Genetic distances, the order of SNP markers, and imputation of missing genotype were calculated by the function *mstmap* in R/ASMap by using the minimum-spanning-tree (MST) algorithm. The Kosambi mapping function was applied during calculation of genetic map distance. The R package LinkageMapView (https://cran.r-project.org/package=LinkageMapView, Ouellette et al. 2018) was used to produce the linkage map and determine the physical position of each SNP marker.

Genotypic data for the SNP markers included in the linkage map were used for QTL analysis combined with the average IS score for each F_3_ family. QTL analysis of IS score was performed in R/qtl. Interval mapping to detect QTLs for resistance to CcM-1 was performed with the nonparametric model (IM-NP) based on the Kruskal–Wallis rank-sum test (Kruglyak and Lander 1995), which is more robust for data deviating from normality. We performed composite interval mapping (CIM, Zeng 1994) to detect QTLs for resistance to 104-T. The parameter settings for CIM were as follows: Haley–Knott regression method and a 1-cM walking speed along chromosomes. The LOD thresholds for QTL detection in IM-NP and CIM were determined by 1,000 permutations. The function *fitqtl* procedure was used to estimate the additive effect, dominant effect, and explained phenotypic variance of each QTL. The genome annotation of the cucumber genome was used to identify the location and predicted function of candidate genes from the 1.5-LOD intervals of significant QTLs. A tool “Display Synteny Blocks” at CuGenDB were used to identify the regions of Gy14 v. 2 reference genome syntenic with the detected QTL regions.

## Results

### Population segregation of disease response to races 0 and 1 of *C. orbiculare*

Analysis of the IS levels of 196 F_2:3_ families for 104-T (race 0) and CcM-1 (race 1) in the cotyledon assay showed a continuous distribution (Fig. 1AB). However, many F_2:3_ lines with a high IS for susceptibility to CcM-1 were segregated, and the distribution peak was at the maximum IS of 7.0 (necrosis of entire cotyledon and seedling wilted; Fig. 1B). This result suggested polygenic control of anthracnose resistance to both 104-T and CcM-1 in this population. The scatter plots of IS levels for each of 104-T and CcM-1 in the cotyledon assay (Fig. 1C) revealed a low correlation (Pearson’s correlation coefficient *r* = 0.47) between resistance to the two strains. Infection with 104-T led to almost no lesions on cotyledons of G100 plants, indicating high resistance, but chlorotic lesions formed on B1 cotyledons, expanding until the entire cotyledon was chlorotic (Fig. 1D). Although F1A and F1B also showed a few small lesions with a few chlorotic lesions, the average IS scores of 1.7 for F1A and 1.6 for F1B suggested that plant response to infection with 104-T was more similar to that of G100 than that of B1 (Fig. 1D). Infection with CcM-1 led lesions on G100 cotyledons typical of HR; necrotic lesions developed rapidly, but the lesion area was limited, and the boundaries of the lesions were not chlorotic (Fig. 1D). In contrast, lesions on B1 and F_1_ cotyledons expanded, had chlorotic boundaries and merged with other necrotic lesions to cover the whole area of the cotyledon (Fig. 1D). Infected B1 seedlings showed not only necrosis on the cotyledons, but the seedlings were wilted. Among the F_2:3_ families, a few lines exhibited lower IS scores than G100, and a few showed IS scores as high as B1, following infection with either 104-T and CcM-1.

**Fig. 1.**
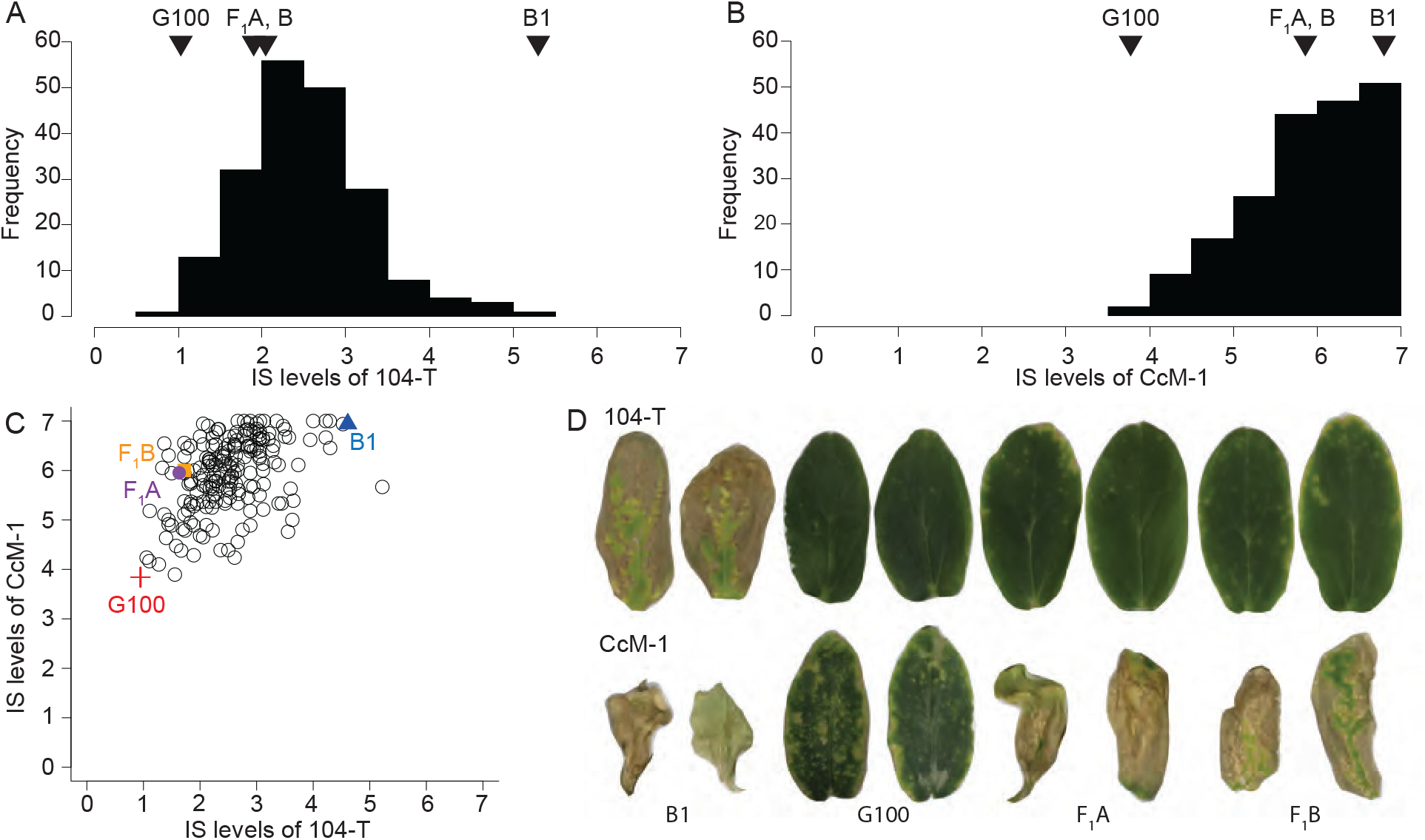
Frequency distribution of the infection severity (IS) levels in 196 F_2:3_ families derived from a B1 × Ban Kyuri (G100) cross assessed by cotyledon assay using the 104-T (A) and CcM-1 (B) strains of *C. orbiculare*. Scatterplot of IS levels following infection with 104-T versus CcM-1 (C), and infected cotyledons of parent lines and F_1_ progenies (D)

The highest five and lowest five F_2:3_ lines in susceptibility to each of 104-T and CcM-1 in the cotyledon assay were selected and tested in the seedling assay. We infected 20 plants from each of selected F_2:3_ lines with different resistance levels, the parents (B1, G100), and the two reciprocal F_1_ lines (F_1_A, F_1_B) with 104-T and CcM-1. Significant positive correlations between IS scores in the cotyledon assay and the seedling assays were observed for both 104-T (*r* = 0.78) and CcM-1 (*r* = 0.83) (Online resource 1A). For plants infected with 104-T, the IS in the cotyledon assay tended to be smaller than those in the seeding assay (Online Resource 1A). For plants infected with CcM-1, the IS in the cotyledon assay tended to be higher than those in the seedling assay, and few seedlings had an IS score of 7.0, suggesting that smaller seedlings were more likely to wilt and die during CcM-1 infection (Online Resource 1A). Plants of G100, B1, F_1_A, and F_1_B, showed very similar symptom severity in both the cotyledon and seedling assays following infection with 104-T and CcM-1 (Online resource 1B).

### Genotyping and linkage map construction

ddRAD-seq analysis using 406 F2 individuals returned an average of 0.2 million high-quality reads per sample, and showed that 27,455 SNPs were polymorphic between the two parental lines. The criteria described in the Materials and Methods were used to filter the SNPs. A total of 955 SNP markers were used for linkage map construction. The linkage map contained seven linkage groups (LGs) and spanned 1572.1 cM, with an average distance between markers of 1.7 cM (Online Resource 5). The linkage map contained long regions (>10 cM) with no markers: 19.5 cM on Chr. 1 and 12.5 cM on Chr. 2 (Online Resource 2, 5).

### QTL mapping of resistance to races 0 and 1 of *C. orbiculare*

Mapping of QTLs for resistance to 104-T and CcM-1, using CIM and IM-NP respectively, produced different results in terms of chromosome position and LOD score, with LOD significance thresholds of 5.99 and 3.53, respectively (Fig. 2, Table 1). For resistance to 104-T, we detected two QTLs: *An5* (LOD: 14.8) on Chr. 5 and *An6*.*2* (9.3) on Chr. 6 (Fig. 2, Table 1). For resistance to CcM-1, we detected four QTLs: *An1*.*1* (LOD: 6.4) and *An1*.*2* (5.8) on Chr. 1, *An*.*2* (8.0) on Chr. 2, and *An6*.*1* (4.6) on Chr. 6 (Fig. 2, Table 1). Previously reported QTLs for resistance to *C. orbiculare* include a major QTL, *cla* (*CsSGR*, Chr. 5), and a minor QTL, *cla3*.*1* (Chr. 3) (Pan et al. 2018). On Chr. 5, *An5* was located at 27.7-28.9 Mb, whereas the reported *cla* (*CsSGR*) was located at 0.2 Mb in Chinese long v3 reference (CuGenDB); furthermore, G100 is reported to be homozygous for the wild-type alleles of *CsSGR* (Matsuo et al. 2022). Consequently, the six QTLs detected here in G100 must be independent novel QTLs for resistance to 104-T and CcM-1.

**Table 1.** QTL anthracnose resistance detected in196 F2:3 families derived from B1 x G100 cross

| Strain | QTL<sup>a</sup> | Chr | QTL Peak<br>Location (cM) | LOD | Additive<br>effect (*a*) | Dominance<br>effect (*d*) | PVE<sup>b</sup> | Total PVE<sup>b</sup> | Range (cM)<sup>c</sup> | Range (Mb)<sup>d</sup> | Range (Mb)<sup>e</sup> | Annotated genes<sup>f</sup> (R genes<sup>g</sup>) |
| --- | --- | --- | --- | --- | --- | --- | --- | --- | --- | --- | --- | --- |
| 104-T | *An5* | 5 | 190.0 | 14.82 | -0.43 | -0.22 | 21.55 | 38.74 | 182.8-196.0 | 27.7-28.9 | 29.7-30.8 | 176 (1) |
| 104-T | *An6.2* | 6 | 212.0 | 9.32 | -0.34 | -0.12 | 9.72 |  | 207.0-214.6 | 28.-28.6 | 29.8-29.9 | 27 (1) |
| CcM-1 | *An1.1* | 1 | 78.5 | 6.37 | -0.15 | -0.09 | 1.75 | 44.39 | 29.1-97.0 | 3.9-7.5 | 3.8-7.5 | 613 (53) |
| CcM-1 | *An1.2* | 1 | 213.0 | 5.78 | -0.24 | -0.19 | 5.13 |  | 197.2-221.0 | 22.48-26.0 | 22.5-26.0 | 410 (8) |
| CcM-1 | *An2* | 2 | 4.3 | 8.00 | -0.46 | 0.86 | 16.96 |  | 0.0-44.89 | 0.5-6.2 | 0.3-6.0 | 690 (32) |
| CcM-1 | *An6.1* | 6 | 174.9 | 4.64 | -0.37 | -0.08 | 9.704 |  | 144.5-197.5 | 23.3-28.2 | 24.3-29.5 | 759 (16) |

<sup>a</sup> QTL names were assigned according to the names of corresponding loci (reviewed by Wang et al. 2020).
<sup>b</sup> Percent of the phenotypic variation explained by the QTL
<sup>c</sup> Genetic distance of the 1.5-LOD interval of the QTL
<sup>d</sup> Physical distance of the 1.5-LOD interval of the QTL in Chinese long v3 reference genome
<sup>e</sup> Syntenic region of the 1.5-LOD interval of the QTL in Gy14 v2 reference genome
<sup>f</sup> Number of potential candidate genes located in 1.5 LOD support interval of QTL
<sup>g</sup> Annotated as R gene like (RLP, RLK, NLR) in 1.5 LOD support interval of QTL

**Fig. 2.**
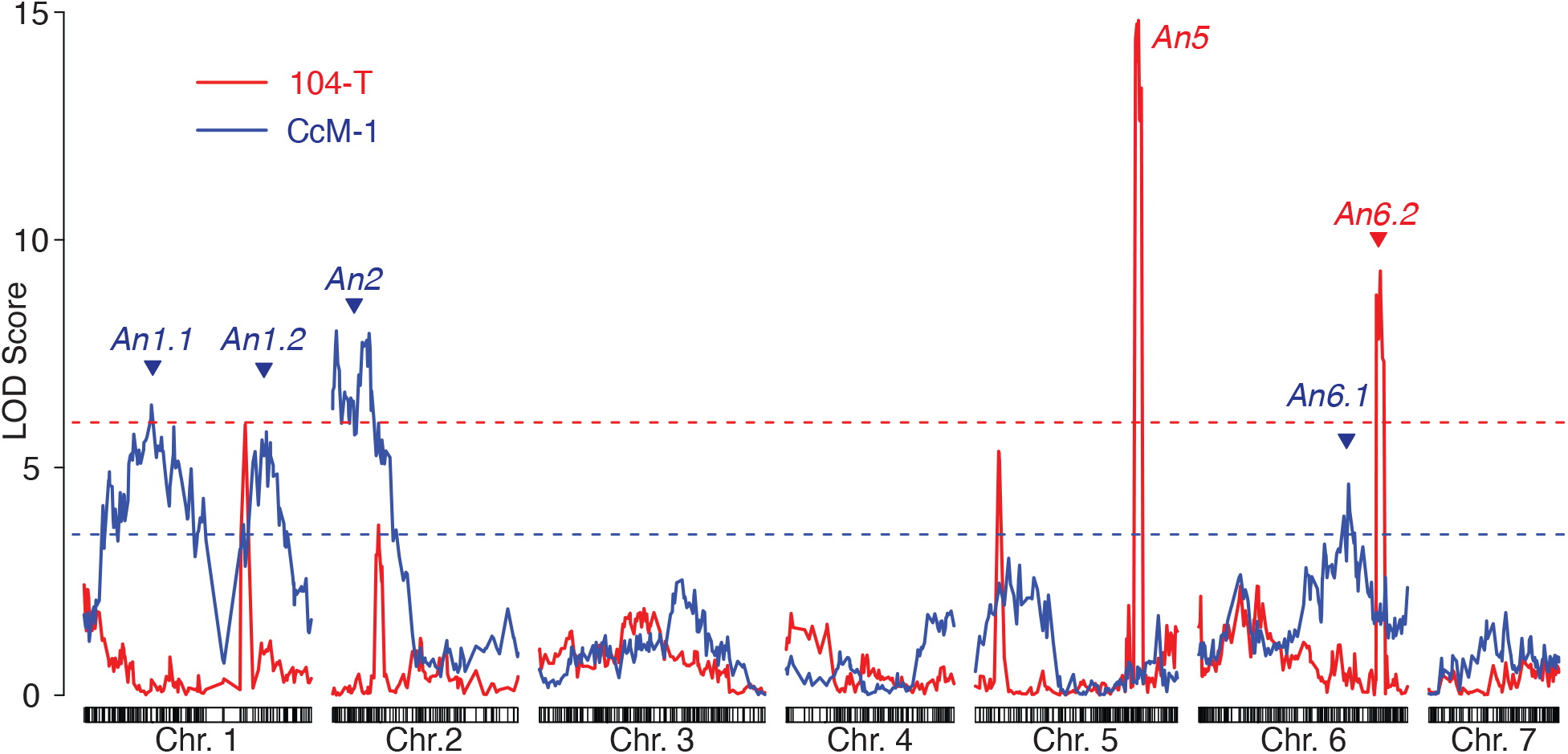
Logarithm of the odds (LOD) score curves showing the positions of each QTL peak for anthracnose resistance detected by CIM (104-T) and IM-NP (CcM-1). Dashed lines show LOD thresholds. We designated QTLs following the nomenclature used by (Wang et al. 2020).

Focusing on the inheritance of the novel QTLs, the absolute value of additive effect (*a*) was higher than that of dominance effect (*d*) (|*a*|>|*d*|) in *An1*.*1, An1*.*2, An5, An6*.*1* and *An6*.*2* (Table 1). However, the absolute value *a* was lower than that of *d* (|*a*|<|*d*|) in *An2* (Table 1). Measuring the effect of QTL genotype by plotting the average IS for each genotype of the SNP closest to the LOD peak of each QTL in the mapping population, for three QTLs on *An1*.*1, An1*.*2* and *An5*, heterozygotes (AB, A: B1 allele; B: G100 allele) had an IS close to that of homozygotes for the G100 allele (BB) (Online resource 3AC). For the three other QTLs, on *An2, An6*.*1* and *An6*.*2*, the average IS were intermediate between the average IS of the AA and the BB genotypes, and each genotype had a linear distribution of IS scores (Online resource 3AC). These results suggested that all the detected QTLs show co-dominant inheritance, but *An2* may show overdominance inheritance. Epistasis and interactions among the detected QTLs were not observed. The total phenotypic variance explained (PVE) by two loci for resistance to 104-T was 38.7%, and *An5* showed a higher PVE (21.6%) than that of *An6*.*2* (9.7%) (Table 1). Thus, *An5* is considered to be a major QTL of G100 resistance to 104-T. By combining two QTLs of BB genotype, a lower IS score indicated the two loci combined to confer strong resistance to 104-T (Online resource 3B).

The 1.5-LOD intervals of the QTLs associated with 104-T resistance were relatively narrow: 11.14 cM for *An5* (182.83 to 195.96 cM) and 7.55 cM for *An6*.*2* (207.04 to 214.59 cM). These intervals corresponded to 1.2 Mb (27.7 to 28.9 Mb) and 0.1 Mb (28.5 to 28.6 Mb) in the cucumber reference genome, respectively, and contained 176 candidate genes for *An5* and 27 for *An6*.*2* (Table 1, Online resource 6). Of known PRRs, only two genes annotated as *lysM domain receptor-like kinase 3* (*CsaV3_5G036150*) and *wall-associated receptor kinase-like* (*CsaV3_6G048820*) are located in the regions of *An5* and *An6*.*2*. These two candidate genes are syntenic with *CsGy5G026460* (*LysM domain receptor-like kinase 3*, Chr. 5: 30381120-30382412) and *CsGy6G032890* (*Wall-associated receptor kinase galacturonan-binding protein*, Chr. 6: 29888530-29891314), respectively, in the Gy14 v. 2 genome.

The PVEs of QTLs for resistance to CcM-1 showed a wide range (1.75–16.96%), and the PVE of *An2* that showed the highest LOD score were also the highest (16.96%) (Table 1). Thus, *An2* is considered to be a major QTL in the G100 resistance to CcM-1. Among the minor QTLs contributing to resistance to CcM-1, *An6*.*1* showed the second highest PVE but the lowest LOD peak (Table 1). However, combination of *An2* and either of minor QTLs did not show evident difference on IS levels, the second most important QTL were not able clearly to be confirmed by 196 F_2:3_ families used for QTL mapping (Online resource 3D). The 1.5-LOD intervals of the QTLs associated with CcM-1 resistance were broad (Table 1). The region of each QTL contained 410-759 genes, including 8–53 PRRs (Table 1, Online resource 6).

## Discussion

We detected six novel QTLs related to resistance to anthracnose (2 QTLs for race 0, strain 104-T and 4 QTLs for race 1, strain CcM-1). Although a previous study suggested that two QTLs (*cla, cla3*.*1*) control resistance to *C. orbiculare* race 1 (Pan et al. 2018), we detected a greater number of QTLs on different positions from above two QTLs, indicating that G100 resistance to race 1 is controlled by different genes from the resistant accession in the previous study. We found wide quantitative variation among F2:3 families, their parental lines, and F_1_ lines in the degree of resistance. As expected, G100 was highly resistant to 104-T and moderately resistant to CcM-1 in the cotyledon and seedling assays. In the cotyledon assay, the IS scores for 104-T and CcM-1 were weakly correlated (Fig. 1B). We did not identify any common QTLs for resistance to races 0 and 1, and resistance to the two races appears to be controlled by independent genes (Online resource 4). Thus, it appears that G100 is the rare material possesses a combination of novel independent resistance QTLs that confer resistance to both race 0 and 1 of *C. orbiculare*.

The cucumber genome contains hot spots where disease-resistance QTLs/genes are clustered (Wang et al. 2020). Comparison with the Gy14 v. 2 reference genome (Table 1) showed that all of the QTLs were detected in these hot spots; in particular, *An5, An6*.*1* and *An6*.*2* map to regions of Chrs. 5 and 6 that contain disease-resistance genes to PM, DM, FOC, gummy stem blight (*Didymella boryniae*) and Cucurbit yellow stunting disorder virus (Online resource 4). In cucumber breeding, positive correlations have been observed between resistance to different pathogens, such as DM and PM; FOC and scab; different potyviruses (Wang et al. 2020). However, G100 does not exhibit resistance to DM, PM, or TLS (data not shown), suggesting that the six QTLs detected in this study are involved in specific resistance to *C. orbiculare* and are not tightly linked to resistance genes to these disease.

Much effort has been devoted to genetic analysis of anthracnose resistance (Abul-Hayja et al. 1978; Barnes and Epps 1952; Busch and Walker 1958; Fanourakis et al. 1987; Linde et al. 1990; Sitterly 1973; Thompson and Jenkins 1985; Wyszogrodzka et al. 1987), to developing molecular markers for selection of resistant plants (Wang et al. 2007; Li et al. 2008), and to identifying a causal gene (*cla, Cssgr*) for anthracnose resistance (Pan et al. 2018; Wang et al. 2018). These studies, except for Wang et al. (2007) and Li et al. (2008), generally used resistant parents that have PI 197087 (an accession from India) and its derivative lines (e.g., Gy14 and WI2757) in their ancestry, and thus the source of the resistance allele (*cla*) in all of them traces back to PI 197087 (Pan et al. 2018). The resistance allele has been widely introgressed into slicer and pickling type cultivars in the USA for over 60 years (Pan et al. 2018; Wang et al. 2018), so it is unlikely there is any linkage drag that would hamper breeding in slicer/pickling cucumber. In contrast, G100 is derived from China, with fruit morphology similar to north Chinese–type oriental cucumbers; it is preserved in the Genebank of the National Agriculture and Food Research Organization, and it is unrelated to PI 197087(Matsuo et al. 2022). The G100 genes for resistance to *C. orbiculare* could be introgressed into other cultivars by using data on SNPs associated with the QTLs detected in this study. However, QTLs associated with fruit morphology and quality in cucumber (Pan et al. 2020; Shimomura et al. 2021; Wang et al. 2020) are located near the resistance QTLs in G100. Therefore, a combination of DNA markers tightly linked to genes for anthracnose resistance and those for fruit morphology/quality will be required to breed for both characteristics.

G100 showed strong resistance to 104-T (race 0), with almost no lesions, and B1 showed severe symptoms, whereas the F_1_ lines showed symptoms similar to those of G100 (Fig. 1C, Online Resource 1B). Hence, the G100 gene for resistance to 104-T must be dominant, given the resistance phenotype of the F_1_ lines and the genotype effect of *An5* and *An6*.*2* in F2:3 families (Table 1, Online resource 4A). *R* genes play a crucial role in plant immune responses, and dominant *R* genes in plants usually encode PRRs that provide full or partial resistance to pathogens (Kourelis and van der Hoorn 2018). Among the PRRs, RLKs/RLPs play a crucial role in PTI and MAMPs-triggered immunity (MTI) which provide partial and sometimes even complete resistance to pathogen (Lacombe et al. 2010; Dubouzet et al. 2011). Combining QTL mapping results (Table 1) with gene annotation of the cucumber genome (Online resource 6) suggests that the causal genes for resistance to 104-T that correspond to QTLs *An5* and *An6*.*2* are *lysM domain receptor-like kinase 3* (*CsaV3_5G036150*) and *wall-associated receptor kinase-like* (*CsaV3_6G048820*), respectively. *LysM receptor-like kinase/proteins* (*LysM-RLK/Ps*), including *lysM domain receptor-like kinase 3*, encode the LysM domain, which can bind chitin-related oligosaccharides (Kaku et al. 2006). In *Arabidopsis, AtCERK1* (Miya et al. 2007), *AtLYK5* (Cao et al. 2014), and *AtLYM2* (Faulkner 2013) are required for chitin sensing, and *AtLYM1* and *AtLYM3* are required for peptide glycan (PGN) sensing (Willmann et al. 2011). RNAi silencing of *OsLYP4* and *OsLYP6* in rice confirmed that these RLPs are involved in chitin and PGN perception (Liu et al. 2013). The *LysM-RLK/Ps* is located on the cell membrane surface, recognizes PAMPs/MAMPs such as chitin and PGN, and plays a critical role in immune function (Wirthmueller et al. 2013; Boutrot and Zipfel 2017).

*Wall-associated protein kinases* (*WAKs*), including *wall-associated receptor kinase-like* (*CsaV3_6G048820*), are a group of RLKs identified in *Arabidopsis* (Kohorn et al. 2009). Some *WAKs* were identified as *R* genes, such as *OsWAK1* (resistance to rice blast disease, *Magnaporthe oryzae*; Li et al. 2009b); *qHSR1/ ZmWAK1* (resistance in maize to head smut, *Sporisarium relianum*; Zuo et al. 2014); and *Htn1/ ZmWAK-RLK1* (resistance in maize to northern corn leaf blight, *Exserohilum turcicum*; Hurni et al. 2015). Further analysis to identify the causal genes of *An5* and *An6*.*2* is needed, and revealing the genetic mechanism of G100 resistance to *C. orbiculare* 104-T might further our understanding of *LysM-RLK/Ps* and *WAKs* associated with race-specific resistance in cucumber. In addition, many virulence-related genes of *C. orbiculare* have been studied, mainly in 104-T, including secreted proteins and effectors involved in PTI (Irieda et al. 2014; Azmi et al. 2018; Kumakura et al. 2019; Chen et al. 2021). Identifying the causal genes of *An5* and *An6*.*2* may lead to identification of 104-T avirulence genes (AVRs) and elucidate the mechanism by which *C. orbiculare* infects its cucurbit host.

QTL analysis revealed that resistance to CcM-1 in G100 appears to be polygenic, controlled by a major QTL, *An2*, and three QTLs, *An1*.*1, An1*.*2*, and *An6*.*1* (Table 1). The 1.5 LOD-support interval for each of *An1*.*1* and *An1*.*2, An2* and *An6*.*1* was wide, and many candidate genes, including NLRs and RLPs/RLKs, are located in these regions (Table 1, Online resource 6). As described in the Introduction, in cucumber NLRs may play a lesser role in cultivar-specific resistance than RLPs/RLKs, such as *CsLRK10L2*, which was identified as a candidate gene for *dm4*.*1*.*2*, one of the sub-QTLs for resistance to DM in PI 197088 (Berg et al. 2020). The causal genes for *An1*.*1, An1*.*2, An2*, and *An6*.*1* that confer resistance to *C. orbiculare* race 1 may also be PRRs. The gene-for-gene theory predicts that most *R* genes are dominant, and have characteristic structures, such as LRR domains (Martin et al. 2003), whereas field resistance genes are recessive, and often have entirely different structures from *R* genes (Fukuoka et al. 2009; Pavan et al. 2010). Based on genotype effects alone, the resistance QTLs to CcM-1 in G100 were dominant or co-dominant or overdominance (Online resource 3B, Table 1). However, entire cotyledons in B1 and F1s were necrosis, but when G100 was infected with CcM-1, it survived with only HR-like lesions (Fig. 1C, Online resource 1B). This result indicates that *An1*.*1, An1*.*2, An2*, and *An6*.*1* may be recessively inherited field resistance genes regardless the resistance activated through PRRs. Thus, *An1*.*1, An1*.*2, An2*, and *An6*.*1* may encode various structures as field resistance genes.

A further noteworthy feature was that the margins of the lesions formed by CcM-1 (race 1) remained green and did not expand (Fig. 1C). In plants, resistance is not only due to innate immunity through PRRs but also to recessive mutations causing loss of function of host-side factors (susceptibility genes, S genes), which are involved in early pathogen invasion, modulation of host defenses, and pathogen sustenance, which is essential for the infection process (van Schie and Takken 2014). The anthracnose resistance gene already identified in cucumber, *Cssgr*, also acts via loss-of-susceptibility mutations (Pan et al. 2018; Wang et al. 2019) and is one example of how *C. orbiculare* infection can be prevented by S gene mutations. Therefore, to identify the causal genes of *An1*.*1* and *An1*.*2, An2, An6*.*2*, comprehensive analysis based on fine mapping is desirable because the causal mutation might be either an *R* gene (PRRs) or an S gene. Our previous work showed that CcM-1 not only belongs to race 1, but is one of the most virulent *C. orbiculare* strains to cucumber in Japan. *Cssgr* is a gene that should be introduced into new resistant cultivars, but this gene alone is insufficient for CcM-1. Hence, understanding resistance to CcM-1 will provide significant advantages to cucumber breeding. It will be necessary to produce other mapping population, such as near-isogenic lines (NILs) with either of the four resistance QTLs from G100, and identify candidate genes by narrowing down the locus region and assessing the effect and heritability of each QTL. Furthermore, pyramiding the G100 resistance QTL with *Cssgr* could provide more robust resistance to anthracnose. Going forward, we need to develop linkage markers that can be used in breeding programs for highly accurate selection, to assess which combination of major QTL *An2* with each minor QTL shows optimal resistance to CcM-1, and to determine the minimum set of QTLs for breeding.

## Supporting information

Online resource 1

Online resource 2

Online resource 3

Online resource 4

Online resource 5

Online resource 6

## Acknowledgements

We thank Z. Shuang, N. Nishi, K. Takano at the University of Tsukuba for technical assistance. Fungal strains and plant genetic resources were kindly provided by NARO Genebank. This work was partly supported by the Ministry of Agriculture, Forestry and Fisheries of the Government of Japan (Collaborative Research Project on Characterization and Evaluation of Plant Genetic Resources for Food and Agriculture (PGRAsia); Grant number: JPJ007117) and Japan Science and Technology Agency (Support for Pioneering Research Initiated by the Next Generation (SPRING); Grant number: JPMJSP2124).

## Statements and Declarations

### Author contribution statement

HM performed the experiments and conducted data analysis. YY designed the study, supervised the experiments, and participated in data analysis. SI and KS performed ddRAD-seq analysis to detect genome-wide SNPs. HM and YY wrote the manuscript with input from the other coauthors. All authors reviewed and approved this submission.

## Data Availability

### Compliance with ethical standards

The experiment conducted complies with the laws of Japan.

## Figure Captions

**Online resource 1** Correlation of F_2:3_ families derived from a B1 x G100 cross assessed by cotyledon and seedling assay using 104-T and CcM-1 (A). Infected seedling of parental lines and F1 progenies by 104-T and CcM-1 (B).

**Online resource 2** Linkage map for B1 x G100 F_2_ population.

**Online resource 3** Effects of single QTL, and combination of major QTL with each minor QTL to 104-T (A), and CcM-1 (B). AA: homozygotes for B1 allele, AB: heterozygoutes for B1 and G100 alleles, BB: homozygotes for G100 allele.

**Online resource 4**. Physical location of *An1*.*1-An6*.*2* compared with resistance genes/QTLs to other disease. This figure is focused on only Chr. 1, 2, 5, and 6, additing *An1*.*1-An6*.*2* based Wang et al. 2020.

