## Supplementary material for "Novel QTLs for cucumber resistance to two *Colletotrichum orbiculare* strains of different pathogenic races": Online resource 1

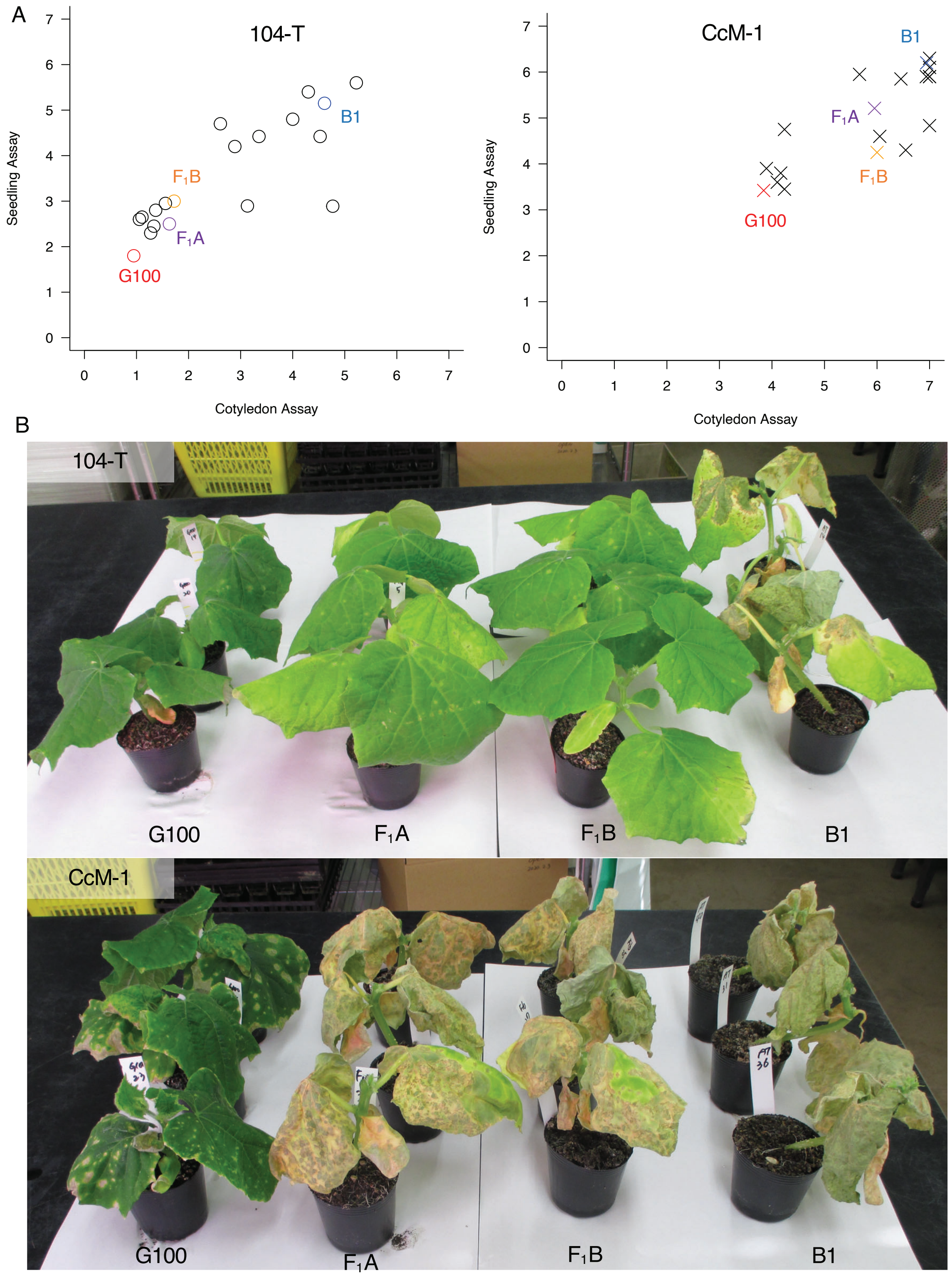


**Online resource 1** Correlation of $F_{2:3}$ families derived from a B1 x G100 cross assessed by cotyledon and seedling assay using 104-T and CcM-1 (A). Infected seedling of parent lines and $F_1$ progenies by 104-T and CcM-1 (B).
