## Supplementary figures and images for "Novel QTLs for cucumber resistance to two *Colletotrichum orbiculare* strains of different pathogenic races"

### Online resource 2

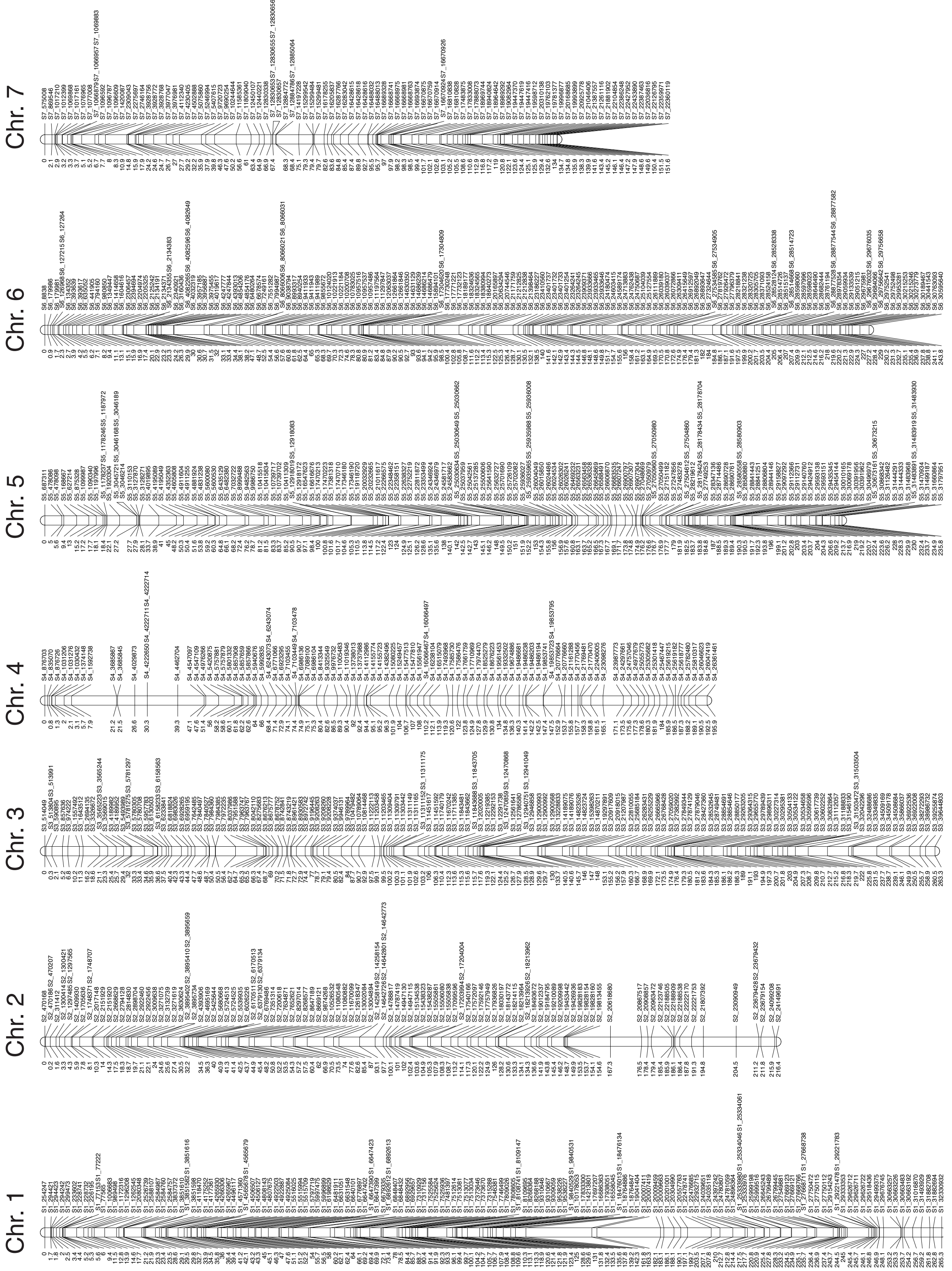


**Online resource 2** Linkage map for B1 x G100 $F_2$ population.
