## Supplementary material for "Novel QTLs for cucumber resistance to two *Colletotrichum orbiculare* strains of different pathogenic races": Online resource 3

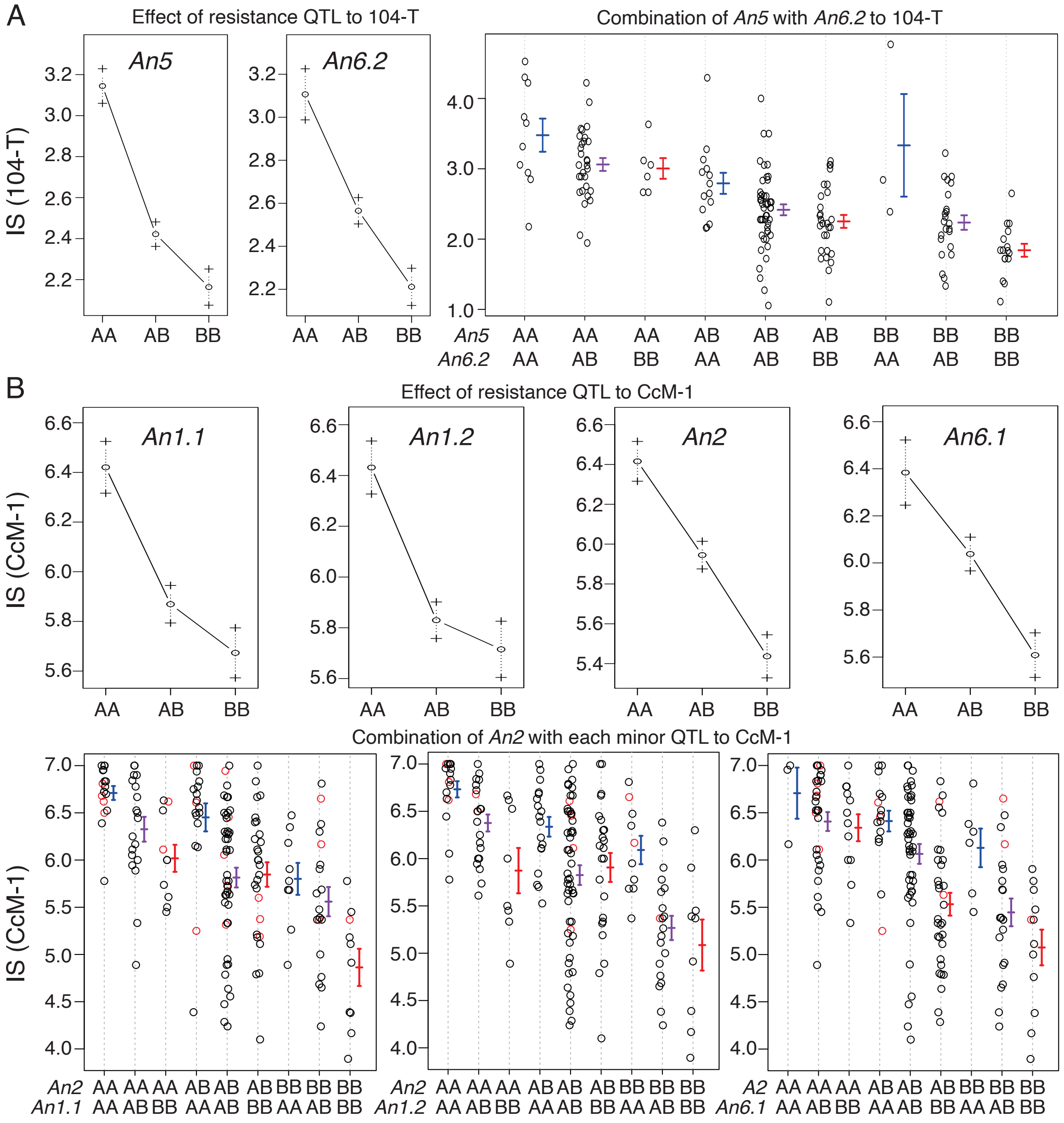


**Online resource 3** Effects of single QTL , and major QTL combination of major QTL with each minor QTL to 104-T (A) and CcM-1 (B). AA: homozygotes for B1 allele, AB: heterozygotes for B1 and G100 alleles, BB: homozygotes for G100 allele.
