## Supplementary material for "Novel QTLs for cucumber resistance to two *Colletotrichum orbiculare* strains of different pathogenic races": Online resource 4

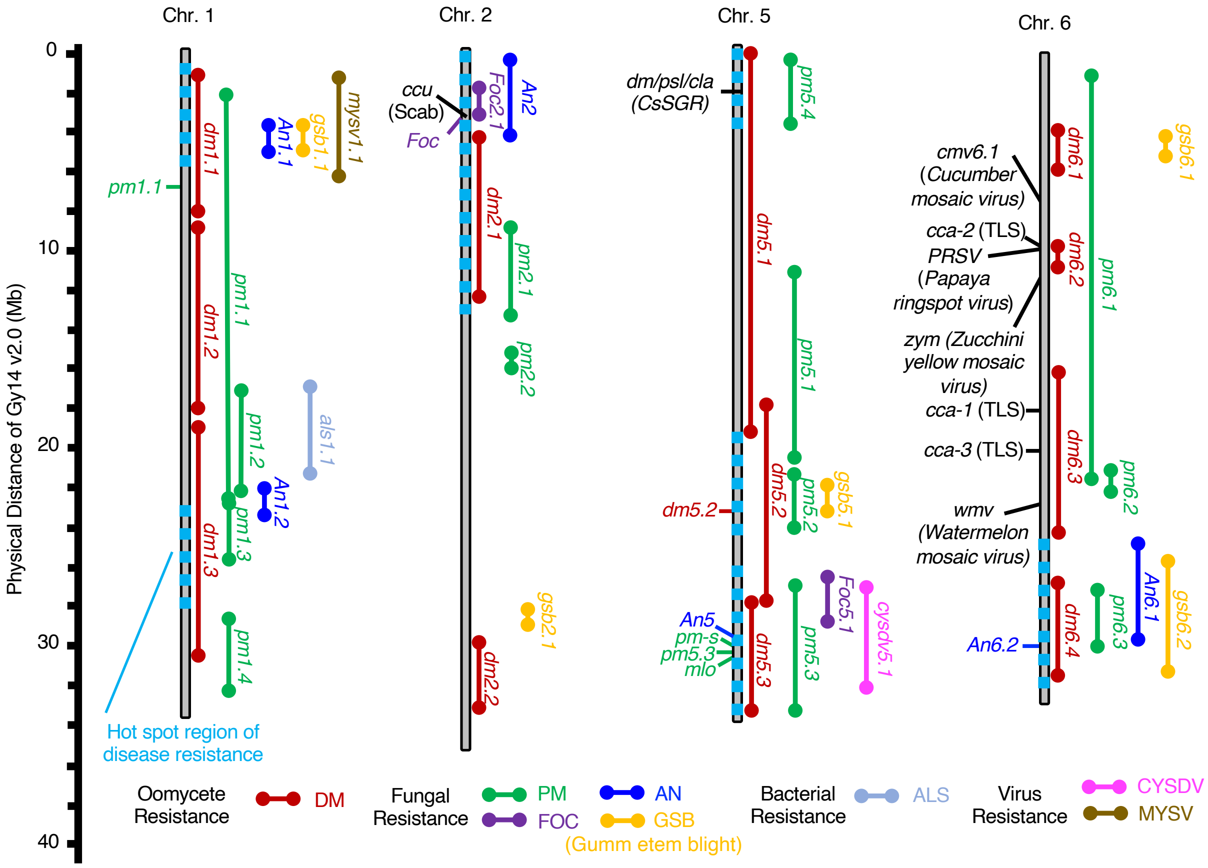


**Online resource 4.** Physical location of *An1.1-An6.2* compared with resistance genes/QTL to other disease. This figure is focused on only Chr. 1, 2, 5 and 6, additing *An1.1-An6.2* based on Wang et al. 2020.
